# Dysphagia as a missing link between post-surgical- and opioid-related pneumonia

**DOI:** 10.1101/2023.10.16.562610

**Authors:** Michael Frazure, Clinton L. Greene, Kimberly E. Iceman, Dena R. Howland, Teresa Pitts

**Affiliations:** Department of Physiology, School of Medicine, University of Louisville, Louisville, Kentucky, United States of America; Department of Neurological Surgery and Kentucky Spinal Cord Injury Research Center, College of Medicine, University of Louisville, Louisville, Kentucky, United States of America; Department of Speech Language and Hearing Sciences and Dalton Cardiovascular Center, University of Missouri, Columbia, Missouri, United States of America

## Abstract

**Objective:** Postoperative pneumonia remains a common complication of surgery, despite increased attention. The purpose of our study was to determine the effects of routine surgery and post-surgical opioid administration on airway protection risk.

**Methods:** Eight healthy adult cats were evaluated for dysphagia in 2 experiments. 1) In 4 female cats airway protection status was tracked following routine abdominal surgery (spay surgery) plus low-dose opioid administration (buprenorphine 0.015mg/kg, IM, q8-12h; *n*=5). 2) Using a cross-over design (2 male, 2 female) cats were treated with moderate (0.02mg/kg) or high (0.04mg/kg) dose buprenorphine (IM, q8-12h; *n*=5) to determine changes in airway protection status or evidence of dysphagia.

**Results:** Airway protection was significantly affected in both experiments, but most severely post-surgically where 75% of the animals exhibited silent aspiration.

**Conclusion:** Oropharyngeal swallow is impaired by the partial mu-opioid receptor agonist buprenorphine, most remarkably in the post-operative setting. These findings have implications for the prevention and management of aspiration pneumonia in vulnerable populations.

## Introduction

Postoperative pneumonia^1–9^ is the third most common side-effect of surgery (regardless of the procedure^10,11^), however there is limited data on the mechanism of this outcome. Two other common risk factors for hospital and community acquired pneumonia are use of opioids^12^ and presence of a swallow disorder (dysphagia)^13^. Swallow is a three-phase behavior with the overall goal of moving food and liquid from the mouth to the stomach ^14,15^. The second phase (pharyngeal) is the riskiest, because the bolus passes by the larynx (entry to the airway) and errors often result in aspiration, which leads to increased risk of morbidity and mortality^16^.

Opioids are commonly prescribed for post-operative pain control, acute or chronic pain management, and/or opioid maintenance therapy ^17–19^. Opioids can depress the gastrointestinal and immune systems^17,20,21^, cough and airway protective reflexes, and have been linked to esophageal dysfunction and aspiration^12,22–25^. Buprenorphine is a commonly prescribed opioid often advertised as safe^26^. It is a partial mu-opioid receptor agonist, which has a lower toxicity profile than full opioid agonists^27,28^. It is widely used for pain management in veterinary practice and is the most prescribed maintenance therapy for human opioid addiction, even in pregnant women^27^.

Opioid receptors are expressed in brainstem regions important for swallow, including pre-motor neurons in the nucleus tractus solitarius, and motoneurons in nucleus ambiguus and hypoglossal motoneuron pools^29,30^. Expanding evidence suggests that the swallow pattern generator is significantly affected by opioid administration^24,31,32^. However, specific effects of buprenorphine and other opioids on oropharyngeal swallow function, and the potential impacts of opioid-induced aspiration, have been the subject of limited study.

In the present study we evaluated the effect of buprenorphine on airway protection in post-surgical animals and healthy non-surgical animals. We hypothesized that buprenorphine would change oropharyngeal swallow function, resulting in airway invasion that may lead to aspiration.

## Materials and Methods

Experiments were approved by the University of Louisville and University of Missouri IACUC and performed on eight healthy adult cats [2 male (5.3±0.4kg) and 6 female (3.4±0.3kg)]. To assess swallow each animal was trained to consume thin liquids and pureed food mixed with 40% barium sulfate (Varibar®, Bracco Diagnostics, Inc.) by volume within a mini c-arm (see Supplemental Methods A; SM-A) as is consistent with videofluoroscopy (i.e., barium/modified barium swallow study) procedures. To assess the impact of a routine abdominal surgery with opioids, and opioids alone, two experiments were performed.

### Experiment 1: Assessment following routine abdominal surgery with low-dose opioid administration

Four adult (>1 year) female cats underwent routine ovariohysterectomy by a veterinarian (CG) according to American Veterinary Medical Association (AMVA) guidelines (see Supplemental Methods B). Animals received pre- and post-operative low doses of buprenorphine hydrochloride (0.015 mg/kg, IM, q8-12h; *n*=5). Swallow was assessed ≥1d prior to surgery, 48-hours post-surgery, and 5-days after the last buprenorphine dose. Assessments were performed 1-hour following buprenorphine administration to ensure peak effect.

### Experiment 2: Assessment with moderate and high-dose opioid administration

Four adult (>1 year) cats (2 female) were randomly assigned to a moderate (0.02mg/kg) or high (0.04 mg/kg) dose of buprenorphine within the feline clinical range (0.01-0.04mg/kg IM). Animals received five doses of buprenorphine (using the post-surgical dosing procedures): One dose administered every 8-12 hours for 48-hours. Swallow assessments were performed before, 24- and 48-hours after the initial dose, and then 24-hours, 72-hours, and 5-days after the last dose. After a one-month washout period, all animals crossed over to the other dose.

### Analysis

Images were exported and analyzed using RadiANT™ DICOM Viewer (Medixant, Poznan, Poland). *Continuous variables* are expressed as means ± standard deviation. SPSS Statistics (v20.0.1.0; IMB®, Armonk, NY) was used to run analysis of variance (ANOVA) and LSD post-hoc tests when appropriate. Differences were considered significant for *p*-values ≤ 0.05. Procedures for measurement of continuous variables are described in Supplemental Methods (SM). *Ratings of safety* were made using an Airway Invasion Scale (AIS, Table 1; SM), and a Timing and Efficiency Scale (Table 1; SM) with two components: 1) Bolus depth at initiation of the pharyngeal phase of swallow (IPS) and 2) Pharyngeal residue of the bolus.

**Table 1.** Ordinal rating scales. The Airway Invasion Scale (AIS) is utilized to track evidence of penetration (yellow box) and aspiration (red box) as well as the amount of material (small vs large). This scale is adapted from Holman and colleagues’ Infant Mammalian Penetration-Aspiration Scale^59^ and Rosenbek’s Penetration-Aspiration Scale^58^. C) The Timing and Efficiency Scale has two components 1) Depth of bolus location at the “Initiation of the Pharyngeal Phase of Swallow” (IPS) and 2) Whether there is “Pharyngeal Residue” following the swallow. These are adapted from the Modified Barium Swallow Impairment Profile (MBSImP) ^75^.

| Score | Airway Invasion Scale (AIS) |
| --- | --- |
| 0 | Material does not enter the airway |
| 1 | Material enters the larynx, remains above the vocal folds, and is ejected from the airway |
| 2 | Material enters the larynx, remains above the vocal folds, and is not ejected from the airway |
| 3 | Material enters the larynx, contacts the vocal folds, and is ejected from the airway |
| 4 | Material enters the larynx, contacts the vocal folds, and is not ejected from the airway |
| 5 | Material enters the larynx, passes below the vocal folds, and is ejected back to the larynx or pharynx |
| 6 | Material enters the larynx, a small amount passes below the vocal folds, and is not ejected despite effort |
| 7 | Material enters the larynx, a large amount passes below the vocal folds, and is not ejected despite effort |
| 8 | Material enters the larynx, a small amount passes below the vocal folds, and no effort is made to eject |
| 9 | Material enters the larynx, a large amount passes below the vocal folds, and no effort is made to eject |
| Score | Timing and Efficiency Scale |
|  | ***Initiation of the pharyngeal phase of swallow (IPS)*** |
| 0 | Bolus head at ramus of mandible |
| 1 | Bolus head in valleculae |
| 2 | Bolus contacts laryngeal surface of epiglottis |
| 3 | Bolus head in pyriform sinus |
| 4 | Bolus spillage to larynx or subglottis |
| 5 | Absent swallow initiation |
|  | ***Pharyngeal residue*** |
| 0 | No residue |
| 1 | Trace residue coating the pharynx |
| 2 | Collection of residue in the pharynx |
| 3 | Majority of bolus remains in the pharynx |
| 4 | Minimal to no pharyngeal clearance |

Ordinal ratings of safety were analyzed using a Wilcoxon signed-rank test.

## Results

### Experiment 1

Figure 1 demonstrates a decline in airway protection during the pharyngeal phase of swallow with 48-hours of post-operative buprenorphine (0.015 mg/kg). Most animals demonstrated aspiration (liquid entering the airway and passing below the vocal folds) without protective response (no cough, throat clear, or cessation of feeding) during post-operative thin liquid feeding. Video 1 (embedded in Fig 1B) demonstrates aspiration during post-operative thin-liquid feeding. Most animals (75%) demonstrated delayed pharyngeal swallow initiation during both thin liquid and puree feeding following post-operative buprenorphine. A Wilcoxon signed-rank test detected a difference in bolus depth at time of initiation of pharyngeal swallow (IPS; Table 1); elevated ratings indicate ingested material spilled deeper into the pharynx or larynx before swallow onset (*z*=-2.3, *p*=0.02). Pharyngeal swallow duration increased by 118±12% during puree feeding (Fig 1), extending the transit time of food through the pharynx, a shared space for breathing and swallow. Maximum esophageal distension increased by 117±5% during puree feeding (Fig 1), with more food accumulating in the esophagus before clearance by primary peristalsis. All animals demonstrated return to functional baseline one-week post-operatively (5-days after last drug administration).

**Figure 1.**
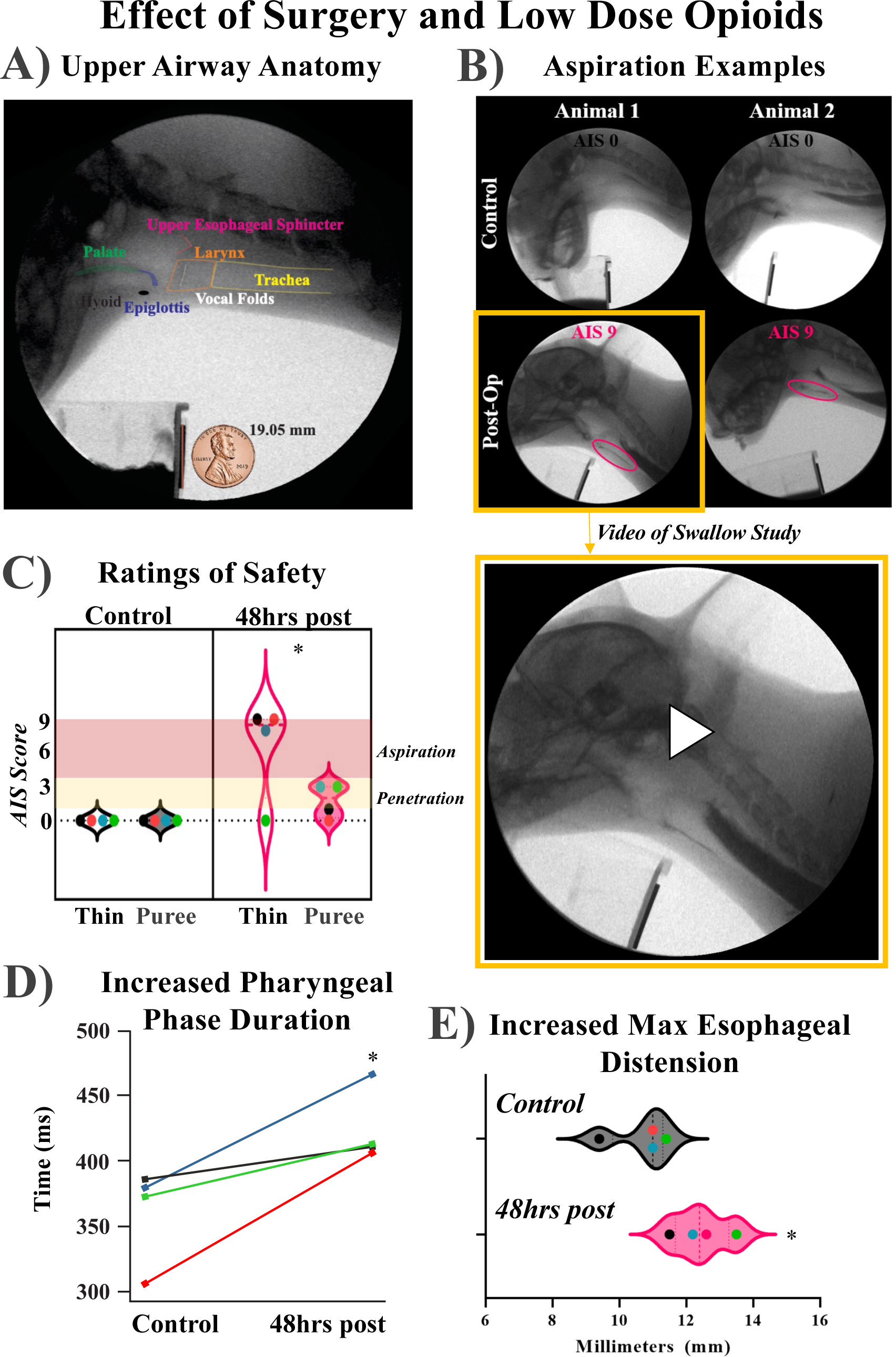
Dysphagia following post-operative low-dose buprenorphine administration. Swallow was evaluated using lateral plane videofluoroscopy. **A)** Anatomic orientation of the oropharynx, pharynx, esophagus, larynx, and trachea. A penny is included for calibration of measurements. Swallowing studies were obtained 48-hours after routine abdominal surgery (ovariohysterectomy), in voluntarily feeding cats 1-hour post intramuscular buprenorphine (0.015 mg/kg) administration. All phases of swallow were impacted. **B)** Three animals presented with significant aspiration without any overt signs or symptom (i.e. silent aspiration). Video 1 (yellow box) demonstrates a feeding bout with increasing amount of contrast agent traveling into the lower airways. **C)** The Airway Invasion Scale (AIS) is an ordinal sale which documents the depth material traveled into the airway during swallow. During control periods all animals-maintained airway protection during puree and thin liquid trials, and following surgery and opioid administration penetration and/or aspiration was present in all animals. A Wilcoxon signed-rank test detected a significant increase in AIS scores (*z* = −2.21, *p* = 0.03). D) Pharyngeal phase duration (onset of hyolaryngeal excursion through upper-esophageal sphincter (UES) closure and return of pharyngeal air space) was significantly longer via [*F*(2, 9)=8.1, *p*=0.01]. and maximum esophageal distension before primary peristalsis [*F*(2, 9)=7.01, *p*=0.015] during puree feeding. E) LSD post-hoc test results revealed that pharyngeal swallow duration was longer during post-operative puree feeding (424±28ms) compared to control (361±37ms). F) Maximum esophageal distension before peristalsis was increased during post-operative puree feeding (12.4±0.9mm) compared to control (10.7± 0.9mm). Swallow metrics returned to functional baseline 1-week post-operatively (5-days after the last dose of buprenorphine). Swallow durations are reported in ms. Distension measures are reported in mm. \**p* < 0.05.

### Experiment 2

Figure 2 demonstrates a decline in airway protection with moderate and high doses of opioids. *24-hours on high dose buprenorphine:* We assessed the effect of opioid administration without surgery on oropharyngeal and esophageal swallow function using two clinical doses and a cross-over design. First, we report the effects of a higher-end clinical dose of buprenorphine (0.04 mg/kg) in a non-surgical setting. One male animal demonstrated total feeding refusal after 24-hours. For the three animals that fed voluntarily, a Wilcoxon signed-rank test detected a difference in IPS depth after 24-hours on buprenorphine than during control (*z*=-2.2, *p*=0.03). ANOVA detected a difference in pharyngeal swallow duration [*F*(2, 7) =27.6, *p*< 0.001]. Post-hoc testing indicated that pharyngeal swallow duration was significantly increased during thin liquid feeding after 24-hours on buprenorphine (121%) compared to control.

**Figure 2.**
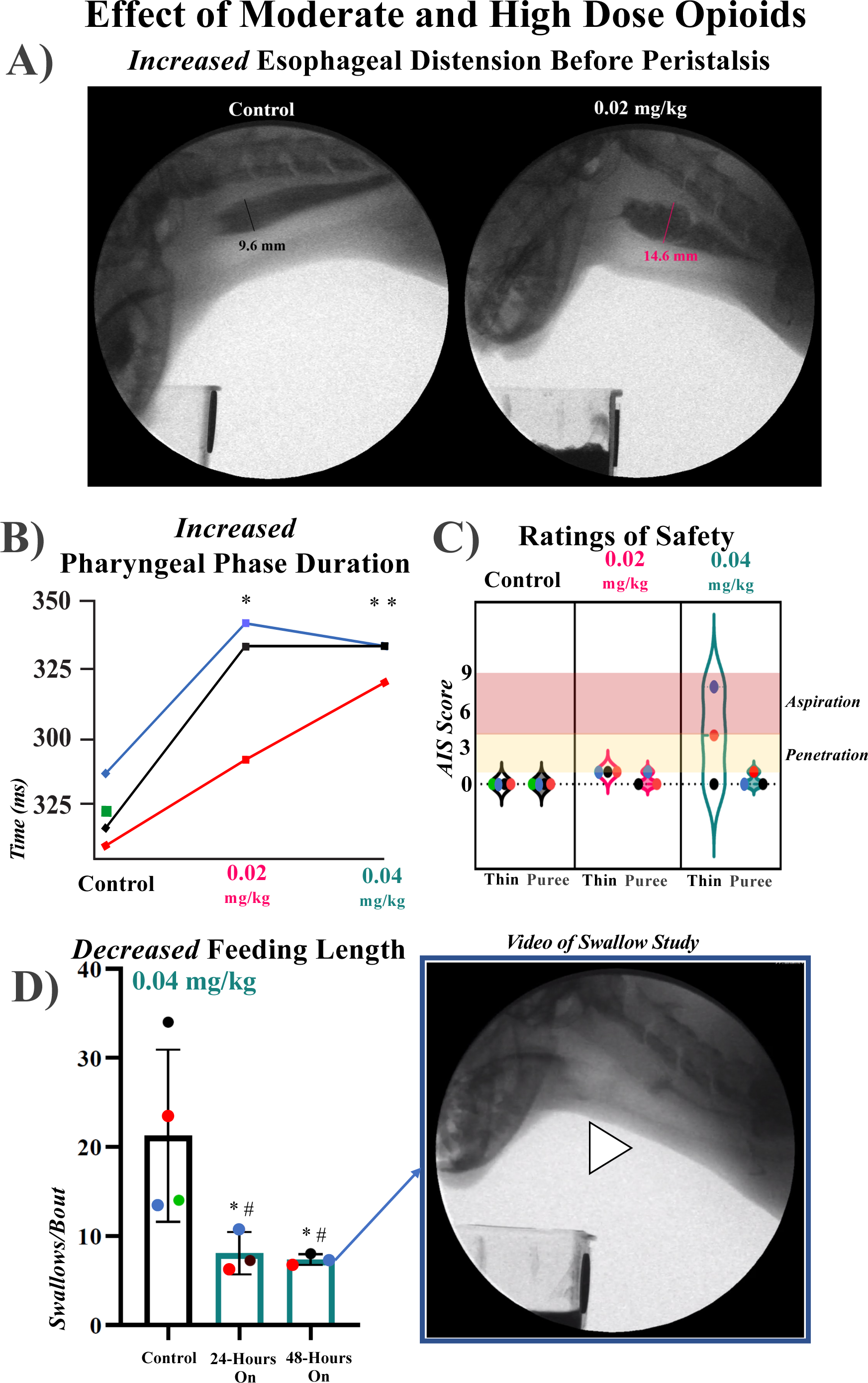
Swallow efficiency is reduced after moderate and high opioid administration. A) Representative videofluoroscopic image depicts increased maximum esophageal distension* after 48-hours on a lower-end clinical dose of buprenorphine (0.02 mg/kg), suggesting decreased sensitivity of esophageal distension reflexes. Maximum esophageal distension before peristalsis increased on average following 48-hours on 0.02 mg/kg (13±2 mm) and 0.04 mg/kg buprenorphine (13±0.8mm) compared to control (12±2mm) but did not reach statistical significance. B) Pharyngeal swallow duration increased during thin liquid feeding after clinical doses of buprenorphine. Analysis of variance (ANOVA) showed differences in pharyngeal swallow duration in the 0.02 mg/kg [*F*(2, 8) = 4.62, *p* = 0.04] and 0.04 mg/kg [*F*(2, 7) = 27.6, *p* < 0.001] data sets. LSD post-hoc test results showed that pharyngeal swallow duration was longer after 48-hours on 0.02 mg/kg (322 ± 26ms) and 0.04 mg/kg buprenorphine (322 ± 13ms) compared to control (271±11ms). C) Feeding bout length decreased during thin liquid feeding after buprenorphine administration. ANOVA showed differences in feeding length in the 0.04 mg/kg data set [*F*(2, 7) = 5.3, *p*=0.04]. LSD post-hoc test results showed that there were fewer swallows per bout after 24-hours (8±2) and 48-hours (7±1) on 0.04 mg/kg buprenorphine compared to control (21 ± 10). One animal demonstrated total feeding refusal at both time points after 0.04 mg/kg buprenorphine. D) A Wilcoxon signed-rank test indicated a change in AIS ratings, with more scores indicative of airway invasion after 48-hours on 0.02 mg/kg buprenorphine (*z* = −2, *p* = 0.05). The mild but significant elevation in AIS ratings indicates that thin liquid laryngeal penetration occurred in all three animals that fed at this time point. AIS ratings were elevated in two of three animals after 48-hours on 0.04 mg/kg buprenorphine, but the effect was non-significant as a group. There was non-transient penetration to the vocal folds in one animal and small volume aspiration in one animal. E) Video 1 shows retrograde flow below the pharyngoesophageal segment, and trace aspiration during thin-liquid feeding in a female animal after 48-hours on 0.04 mg/kg buprenorphine. Of note: \**p* < 0.05, \*\**p* < 0.01, # indicates feeding refusal.

*48-hours on high dose buprenorphine:* One male animal demonstrated total feeding refusal after 48-hours on 0.04 mg/kg buprenorphine. For the three animals that fed voluntarily, AIS ratings were elevated in two of three animals, but the effect was not significant as a group (Fig 2). A Wilcoxon signed-rank test detected a difference in bolus depth at the IPS; IPS ratings were significantly higher following 24-hours on buprenorphine than during control (*z*=-2.2, *p*=0.03). Pharyngeal swallow duration was longer during thin liquid feeding after 48-hours on buprenorphine than during control, extending the time of liquid transit through the pharynx (Fig 2). Video 2 is a representative example of thin-liquid feeding after 48-hours on 0.04 mg/kg buprenorphine in a female animal. All animals resumed voluntary feeding within 24-hours of the last dose of buprenorphine and returned to functional baseline within 5-days of the last dose.

*24-hours on moderate dose buprenorphine:* The moderate dose of buprenorphine (0.02 mg/kg) also had a significant impact on airway protection. After 24-hours AIS ratings elevated, but were just below significance (*z*=-1.9, *p*=0.059). There was laryngeal penetration during thin liquid feeding in 3/4 animals. A Wilcoxon signed-rank test detected a difference in bolus depth at the time of IPS after 24-hours on buprenorphine compared to control (*z*=-2.7, *p*=0.008). An ANOVA showed a difference in swallow duration [*F*(2,8)= 4.62, *p*=0.04] and pharyngeal distension [*F*(2,9)=4.1, *p*=0.05]. LSD post-hoc testing revealed that pharyngeal swallow duration was longer during thin liquid feeding after 24-hours on buprenorphine (325±37) than during control (271±11). Post-hoc testing revealed that pharyngeal distension before swallow increased ∼120% after 24-hours on buprenorphine.

*48-hours on moderate dose buprenorphine:* The same male animal also demonstrated total feeding refusal after 48-hours of moderate dose buprenorphine. The following results pertain to the three animals that fed voluntarily after 48-hours of opioid administration. Figure 2 demonstrates elevated AIS ratings during thin liquid feeding, with laryngeal penetration in all animals. A Wilcoxon signed-rank test detected a difference in bolus depth at the time of IPS; IPS ratings were higher after 48-hours on buprenorphine than during control (*z*=-2.2, *p*=0.03).

Pharyngeal swallow duration increased during thin liquid feeding after 48-hours on buprenorphine compared to control (Fig 2). Post-hoc testing revealed a trend toward increase *(p*=0.07) in maximum pharyngeal distension during puree feeding after 48-hours on buprenorphine (12±0.9) compared to control (11±1). All animals resumed voluntary feeding within 24-hours of the last dose of buprenorphine and returned to functional baseline within 5-days of the last dose.

Inter-rater reliability^33^ was strong [Kappa=0.97 (*p* < 0.001), 95% CI (0.94, 0.98)]. The average two-way random intraclass coefficient (ICC)^34^ for MF was ICC = 0.99 [*F*(32, 32) = 625.9, *p* < 0.001], 95% CI (0.99, 0.99) and for TP was ICC = 0.97 [*F*(32, 32) = 67.2, *p* < 0.001], 95% CI (0.94, 0.99).

## Discussion

This is the first demonstration of dysphagia following routine abdominal surgery with opioid treatment in otherwise young, healthy, adult animals. Further, dysphagia was present after opioid treatment alone, including cessation of feeding after moderate and high doses of buprenorphine in male and female animals.

Opioids are best known for their analgesic effects and abuse potential, but respiratory and gastrointestinal depression are recognized side effects ^35^. Most clinically relevant effects occur via the mu-opioid receptor ^36,37^. Buprenorphine is considered a safe alternative for post-operative pain management and opioid maintenance therapy because it is a partial agonist with a reported ceiling effect at high doses ^27,28^. However, hospitalizations, poison control cases, and potential for lethal interactions with depressants (e.g., benzodiazepines) suggest possible adverse effects ^35^. Full mu-agonists (e.g., morphine) are known to gastrointestinal motility, cough, and related airway protective behaviors, but there is limited information on partial mu-agonists (e.g., buprenorphine) ^17,20,25,38^.

The medullary swallow pattern generator consists of the nucleus tractus solitarius (NTS), nucleus ambiguus (NA), reticular formation (RF) and various cranial and spinal nerve motor nuclei and receives dense inputs ^39,40^. Opioids can disrupt respiratory, cough, emetic, and spinal locomotor pattern generators ^41–44^. Because of this, and the density of the mu-opioid receptors in the NTS and NA, it is unsurprising that opioids disrupt swallow function ^29,45^, however the extent of impact was not predicted at the start of these studies.

Instrumentation is required to definitively evaluate airway protection during swallow ^46,47^. During VFSS, the lateral fluoroscopic view includes the oral cavity, pharynx, larynx, trachea, and proximal esophagus, enabling real-time assessment of airway protection, kinematics, and bolus transit ^48,49^. Because VFSS does not require insertion of a scope and can visualize all phases of swallow, it is an ideal method for animal studies ^50,51^. Translational models of VFSS have been described in the mouse, rat, dog, and neonate pig ^51–54^. While electrophysiological studies of airway protective behaviors in anesthetized cats have been foundational to models of neural control in humans, swallow studies in awake, voluntarily feeding cats have been limited ^40,55,56^. Adult cats have similar mechanisms of airway protection to humans and can feed freely in a fluoroscopic field ^55,57^. As such, a cat model of VFSS during normal and disordered swallow is highly translatable and enables study of central mechanisms using injury or pharmacology not possible in humans ^50^.

We adapted our Airway Invasion Scale (AIS) from clinical and animal-model scales ^58,59^. Because we found that some instances of aspiration involved large amounts of liquid while others involved trace amounts of liquid in the airway, we developed ratings that account for volume. The AIS will permit evaluation of swallow function and therapeutic outcomes in several translational disease models.

Airway invasion was deeper after opioids during thin liquid and puree feeding in both non-surgical groups and post-operative groups, and it occurred without protective response. This is understandable, as mu-opioid agonists suppress cough, and cough is a major protective response to aspiration ^57,60–62^. Importantly, pharyngeal swallow function also declined, resulting in increased airway invasion and depressed protective reflexes. We speculate that opioid-induced oropharyngeal dysphagia is a) under-detected due to suppression of overt indicators of aspiration (e.g., cough) and b) a contributing factor to post-operative aspiration pneumonia.

The most severe aspiration occurred during thin liquid feeding following post-operative buprenorphine administration. After trauma or elective surgery, the stress/inflammatory response is an adaptive process in which cardiometabolic, neuroendocrine, and immune responses are mounted to protect immediate survival ^63,64^. Inflammatory markers peak two days post-injury and return to baseline after one week ^65^. Animals in our post-operative group demonstrated severe aspiration two days after surgery, with return to baseline function after one week. We speculate that post-operative inflammation may have contributed to that severe dysphagia. Moreover, alteration of spinal feedback related to post-surgical changes in abdominal pressures may have disrupted swallow function in the post-operative group ^66–69^.

Swallow initiation, swallow duration, and distension reflexes changed following buprenorphine administration in both non-surgical and post-operative groups. Most animals demonstrated delayed pharyngeal swallow initiation. Bolus location at time of pharyngeal swallow initiation spans from the oral tongue to the pyriform sinus during normal feeding and is variable by consistency and at the population level. Because the glottis is open at rest, there is increased likelihood of airway invasion when a bolus spills to the pharyngeal cavity prior to swallow initiation. Risk of airway invasion with posterior bolus spillage is further increased in the setting of sensory or motor impairment. Our results show that increased depth of food/liquid spillage and delayed onset of pharyngeal swallow resulted in increased airway invasion before swallow following opioid administration.

For bolus transport to occur without airway aspiration, the laryngeal vestibule must remain tightly closed for the duration of pharyngeal swallow. Increased duration of pharyngeal swallow after opioid administration indicates decreased oropharyngeal efficiency, extending the period for airway invasion opportunity.

Pharyngeal distension prior to swallow initiation increased in the non-surgical group, and esophageal distension prior to primary peristalsis increased in the post-operative group. This suggests that opioids decrease the sensitivity of pharyngeal and esophageal distension reflexes, consistent with reports that esophageal circuitry is enriched with mu-opioid receptors, and human studies demonstrating disordered peristalsis and achalasia following opioid administration ^22,23,70–73^. We speculate that increased filling of the pharynx and esophagus increases the likelihood of airway invasion and laryngopharyngeal reflux.

Number of swallows per feeding bout was reduced following buprenorphine without surgery, but unchanged after post-operative buprenorphine. Animals in the non-surgical group required increased direction and fed for shorter periods of time, and one animal demonstrated total feeding refusal. Reduced intake and food refusal are clinical indicators of dysphagia. We propose two explanations for this phenomenon: A) Reduced feeding length was a compensatory mechanism in the non-surgical group, who demonstrated less severe aspiration than the post-operative group and b) poor feeding was a function of higher clinical doses of buprenorphine administered to the non-surgical group.

## Conclusion

Oropharyngeal swallow function is impaired by the opioid buprenorphine, a drug that is considered safe for pain management and treatment of opioid addiction disorder ^19,74^. We propose that risk for dysphagia-related chest infection following opioid administration is greatest in the post-operative setting. We hypothesize that decline in swallow function following opioid administration occurs due to central disturbance of the swallow motor pattern, and depression of aerodigestive sensation and protective reflexes. Our findings have implications for prevention of aspiration pneumonia in post-surgical cases. In the event of post-operative opioid administration, instrumental evaluation of swallowing is indicated for patients at increased risk for dysphagia-related chest infection, including persons who are immunocompromised or aged.

## Supporting information

Supplemental Methods and Results

