## Supplemental Methods and Results for "Dysphagia as a missing link between post-surgical- and opioid-related pneumonia"

**Supplemental Methods A.** Acclimation and Videofluoroscopy

Acclimation is a multi-step process which takes an average of 30 days. Animals were initially acclimated to the examiner during training sessions (< 15 minutes) 3-5x/day. They were then acclimated to transport and temporary housing in our fluoroscopy suite. Animals then progressed from feeding in a deluxe crate ("The Other Door", 42"L x 28"W x 28"H, Pet Gear Inc.; Vermont Juvenile Furniture Manufacturing, Inc, West Rutland, VT), to feeding freely in a natural stance without direction in the fluoroscopic field, and were considered trained following 5 consecutive days of feeding during recording of lateral plane VFSS (30 frames/sec; Fluoroscan® Insight mini-C-arm; Hologic®, Marlborough, MA). Two consistencies were used for each assessment (thin and puree). The thin liquid consistency was water shaken with human-grade canned tuna fish and strained, and the puree consistency was Friskies® branded paté cat food "Turkey and Giblets Dinner" (Purina®, Neenah, WI) both mixed with a commercially available barium sulfate (Varibar®, Bracco Diagnostics, Inc., Milan, Italy) at 40% by volume.

**Supplemental Methods B.** Spay Procedure

Four healthy adult female cats underwent routine ovariohysterectomy performed by a veterinarian specialized in Canine/Feline Practice as certified by the American Board of Veterinary Practitioners (ABVP). Animals received pre-anesthetic treatment with buprenorphine hydrochloride (0.015 mg/kg), acepromazine (0.015 mg/kg) and atropine sulfate (0.015 mg/kg), combined and administered intramuscularly (IM). Anesthesia was induced using isoflurane in a ventilated induction chamber administered via precision vaporizer (5% isoflurane, 5 l/min $O_2$). Once a light anesthetic plane was achieved, animals were removed from the induction chamber, positioned sternally, received eye lubrication, and switched to a facemask for continued anesthesia delivery (5% isoflurane, 1 l/min $O_2$). When a deeper anesthetic plane was achieved, animals were intubated with a 3.5 mm cuffed endotracheal tube for maintenance of anesthesia (2-3% isoflurane, 1 l/min $O_2$) and a 22g intravenous (IV) catheter placed in the cephalic vein. Animals were then repositioned ventrodorsally, and the abdomen shaved and sterilely prepped and draped for surgery. A surgical monitor was used to continuously track pulse, electrocardiogram (ECG), oxygen saturation ($SpO_2$), and end-tidal carbon dioxide ($ETCO_2$), as well as automated, intermittent, non-invasive blood pressure (NIBP) measurements.

Surgical anesthetic depth was confirmed using reflex responses and eye position. A standard midline laparotomy approach (2-3 cm) was performed approximately one inch below the umbilicus and a spay hook used to assist in exteriorizing the left ovary and uterine horn. The suspensory ligament and associated vessels were double clamped with paired transfixion ligatures placed using 3-0 poliglecaprone 25 monofilament suture. The ligament was transected, and the stump inspected for bleeding prior to placement back into the peritoneal cavity. The procedure was then repeated on the right ovary and uterine horn. The broad ligaments were digitally broken down, and the uterine body was exteriorized and similarly double clamped at the cervix, double ligated, transected, and inspected for bleeding. The ovaries and attached uterus were removed and the uterine stump replaced into the peritoneal cavity. Routine closure in three layers (linea and subcutis – continuous; skin - subcuticular) was done and tissue glue (octyl/butyl cyanoacrylate) applied.

After completion of surgery, animals were administered cefazolin sodium (20 mg/kg, slow IV), taken off anesthesia and extubated following return of the pharyngeal swallow reflex. The IV catheter was removed, and patient monitoring continued until recovery was complete before returning to housing. Mean time under anesthesia was 1.3 ± 0.5 hours, mean length of intubation was 1.1 ± 0.5 hours, and mean length of surgery was 0.8 ± 0.4 hours. Buprenorphine (0.015 mg/kg, IM, q8-12h) was administered for post-operative pain control.

**Supplemental Methods C.** Continuous Variables

Images were viewed and analyzed using RadiANT DICOM Viewer (Medixant; Poznan, Poland). Pharyngeal and esophageal distension measures were made using the digital imaging and communications in medicine (DICOM) viewer's length measurement tool. Bolus area was measured using the DICOM viewer's closed polygon measurement tool. Each metric is defined in Supplemental Table 1.

**Supplemental Table 1. Definitions of VFSS metrics.**

| Metric | Definition |
|---|---|
| Oral Phase Duration (ms) | Time from beginning of lapping behavior to pharyngeal swallow initiation |
| Pharyngeal Phase Duration (ms) | Time from onset of hyolaryngeal excursion to UES closure and return of pharyngeal air space |
| Oral to Pharyngeal Phase Ratio | Number of tongue laps prior to pharyngeal swallow initiation |
| Pharyngeal to Esophageal Phase Ratio | Number of pharyngeal swallows prior to initiation of primary peristalsis |
| Total Swallow Count | Number of swallows per feeding bout |
| Pharyngeal Distension (mm) | Bolus width from tongue base to epiglottic rim |
| Esophageal Distension (mm) | Width of food-filled, proximal esophagus prior to initiation of primary peristalsis |
| Bolus Area ($mm^2$) | Area of bolus prior to pharyngeal swallow initiation |

**Supplemental Methods D.** Ordinal Scales

We adapted a categorical Airway Invasion Scale (AIS, Table 1) from Rosenbek and colleagues' 8-Point penetration aspiration scale[58] and Holman and colleagues' infant mammalian penetration-aspiration scale[59]. Novel ratings that reflect volume of aspiration were added to our AIS. A Timing and Efficiency Scale (Table 1) was adapted from Martin-Harris and colleagues' Modified Barium Swallow Impairment Profile (MBSImP)[75]. Two of the 17 physiologic components of swallow described in the MBSImP may be applied to lateral plane videofluoroscopy in the cat: Initiation of the pharyngeal phase of swallow (IPS) and pharyngeal residue. A novel rating that reflects bolus spillage to the upper airway prior to swallow initiation was added to our Timing and Efficiency Scale (Table 1).

**Supplemental Results A.** Reliability and Blinding

Airway Invasion Scale (AIS) scores were made by two speech-language pathologists with the certificate of clinical competence (CCC) from the American Speech Language and Hearing Association (ASHA) and at least five years of experience (MF, TP). Raters scored de-identified images for: Control condition, buprenorphine without surgery, and post-operative buprenorphine.

An inter-rater reliability analysis was performed using Cohen's kappa coefficient to determine consistency among raters [33]. 30% of videos were repeated at random to allow for assessment of intra-rater reliability, determined by a two-way random intraclass correlation coefficient[34]. Reliability statistics were performed using SPSS software (IBM; Chicago IL, USA). Both intra- and inter-rater reliability were considered adequate if equal to or greater than 80%. Ratings of timing and efficiency were made by an ASHA certified speech pathologist (MF) using the same de-identified data set following demonstration of inter- and intra-rater reliability greater than 80%.

The inter-rater reliability for the raters was found to be Kappa = 0.97 ($p < 0.001$), 95% CI (0.94, 0.98). The average two-way random intraclass coefficient (ICC) for MF was ICC = 0.99 [$F(32, 32) = 625.9$, $p < 0.001$], 95% CI (0.99, 0.99). The average two-way random intraclass coefficient for TP was ICC = 0.97 [$F(32, 32) = 67.2$, $p < 0.001$], 95% CI (0.94, 0.99).
